# Integrative reanalysis of public intracellular and vaccine-challenge transcriptomes, informed by single-cell and metabolomic atlases, refines antigen prioritisation for next-generation veterinary *Brucella* vaccines

**DOI:** 10.64898/2026.10.19.760311

**Authors:** Yunwang Zhao, Hongxian Cheng, Shiyi Zhou, Lei He, Chenlin Xiu, Baihai Huang, Xiongxiong Lu

## Abstract

Brucellosis is the most widespread bacterial zoonosis and a leading cause of abortion and infertility in cattle, sheep, goats and pigs. The live vaccines licensed for livestock (*B. abortus* S19 and RB51, *B. melitensis* Rev.1) give incomplete protection, retain residual virulence and induce antibodies that are indistinguishable from those of field infection, and no human vaccine is licensed. Antigen discovery for safer subunit and mRNA alternatives has proceeded one candidate at a time, so it remains unclear which immunogenic proteins are abundant in the intracellular niche that protective immunity must recognise. We reanalysed four public data layers: dual RNA-sequencing of intracellular *B. abortus* and infected murine macrophages (GSE100914), splenic transcriptomes from vaccinated and challenged mice (GSE254248), and single-cell (PRJCA054635) and metabolomic (ST003696, ST003692) atlases. Intracellular residence induced 704 genes and repressed 249, enriching translation functions 2.9-fold. Omp28/BP26 was the second most abundant intracellular transcript (35,652 TPM; log2FC 4.25), whereas the lipoproteins omp16, omp10 and omp19 were 116-637-fold less abundant. Vaccination left a germinal-centre and TCF1+ memory imprint rather than an effector-cytokine signature. A composite score combining abundance, induction and surface exposure reorders the candidate panel, placing Omp28/BP26 and omp2b above current candidates and defining concrete priorities for veterinary vaccine development.

**Simple Summary:** Brucellosis is a bacterial infection that causes abortions, stillbirths and infertility in cattle, sheep, goats and pigs, and it can also pass to people through unpasteurised milk or direct contact with infected animals. Vaccination is the main control measure, but the live vaccines used in livestock do not fully protect animals and make vaccinated animals look infected in the blood tests on which eradication programmes depend. Safer subunit and mRNA vaccines therefore need better target proteins, yet candidates have usually been chosen one at a time from laboratory-grown bacteria, which may not reflect what the bacteria produce inside the animal’s own cells. We reanalysed publicly available datasets that measure gene activity in infected cells and in vaccinated animals. The proteins currently favoured by vaccine developers turn out to be scarce inside host cells, whereas two surface proteins are abundant and strongly switched on. Our ranking, together with the immune features that distinguish protected animals, gives concrete priorities for the next generation of veterinary brucellosis vaccines.

## 1. Introduction

Brucellosis is a globally distributed zoonosis caused by facultatively intracellular bacteria of the genus *Brucella*. Human disease typically presents as undulant fever, but focal complications including spondylitis, endocarditis, neurobrucellosis and orchitis occur in a substantial minority of patients, and up to 30% develop chronic or relapsing illness [1,2]. In livestock the organism causes abortion and infertility, generating severe economic losses and sustaining the reservoir from which human infection is acquired through unpasteurised dairy products or occupational contact [3]. The global burden is estimated at 500,000 new human cases annually, with the highest incidence in the Middle East, Central Asia, Sub-Saharan Africa, Latin America and parts of China [4].

Vaccination is the only realistic route to durable control, yet the field has been constrained by the same fundamental problem for decades. The vaccines licensed for livestock-*B. abortus* S19 and RB51, and *B. melitensis* Rev.1-are live attenuated strains that provide incomplete protection, can induce abortion in pregnant animals, are pathogenic to humans, and provoke antibodies that are indistinguishable from those generated by field infection, thereby destroying the diagnostic basis of eradication programmes [5,6]. For humans, no vaccine is licensed at all. The search for safer subunit, DNA, vectored or mRNA alternatives has consequently produced a large and growing catalogue of candidate antigens, including the outer-membrane proteins Omp25, Omp31, Omp19, Omp16 and Omp10, the periplasmic protein BP26, the ribosomal protein L7/L12, superoxide dismutase, lumazine synthase, malate dehydrogenase and the T4SS effector VirB proteins [7], alongside a growing wave of in silico and mRNA vaccine designs in diverse bacterial pathogens [8-10].

This catalogue has been assembled almost entirely through hypothesis-driven, one-antigen-at-a-time screening, guided by immunodominance in natural infection or by convenient recombinant expression. That approach has an important and under-appreciated blind spot. *Brucella* is an intracellular pathogen that survives within macrophages and dendritic cells by actively remodelling its own transcriptome and by subverting host cell-autonomous immunity [11,12]. The relevant target of a protective response is therefore not the antigen profile of the bacterium as it exists in laboratory culture, but the profile it presents inside the host cell. Whether the established candidates are actually abundant in that niche-and whether highly expressed alternatives have simply been overlooked-has never been systematically assessed.

Two developments now make such an assessment feasible. First, immunoinformatics and reverse vaccinology have matured into practical tools for antigenicity, allergenicity, epitope and structural prediction, and have been applied successfully to *Brucella* and to other intracellular pathogens [13-15]. Second, and more decisively, the public sequence archive has accumulated enough *Brucella*-specific transcriptomic, single-cell and metabolomic data that a genuinely data-driven analysis can now be performed without generating new data. We reasoned that mining this existing record could test a simple, falsifiable hypothesis: that the antigens currently prioritised for *Brucella* vaccine development are not the antigens most abundantly and most strongly induced in the intracellular niche, and that correcting this mismatch would immediately yield better candidates.

We therefore assembled four independent public data layers covering the pathogen transcriptome inside macrophages, the host splenic response to vaccination and challenge, the single-cell composition of the infected spleen, and the Brucella metabolome, reanalysing the two transcriptomic layers in depth and using the single-cell and metabolomic atlases for biological context (Section 3.6). We show that the intracellular transcriptome is dominated by a small number of highly expressed surface-associated proteins whose ranking diverges substantially from the current development pipeline, and we propose a transparent composite score for prioritising candidates. We further show that protective vaccination leaves a distinctive germinal-centre and memory-lymphocyte imprint, and we define a prospective spatial transcriptomic experiment to determine where in the splenic architecture that imprint is functionally anchored. Throughout, we have been explicit about the limits of publicly available data, including the complete absence of spatial transcriptomic data for *Brucella* and the confounding tissue-composition effects in bulk spleen RNA-sequencing, so that the conclusions can be weighed appropriately.

## 2. Materials and Methods

### 2.1. Data Sources and Accession Numbers

No new experiments were performed. All datasets analysed here were obtained from public repositories and are individually identified below. The complete set of accessions, sample compositions and analytical roles is summarised in Table 1.

**Table 1.** Public datasets integrated in this study. All data were obtained from open repositories and reanalysed as described in Section 2.2.

| Accession | Repository | Host / organism | Design and content | Role in this study |
| --- | --- | --- | --- | --- |
| GSE100914 | NCBI GEO | *B. abortus* + Mus musculus | Dual RNA-seq: intracellular *Brucella* (24 h post-infection of BMM) versus free-living bacteria; parallel host BMM transcriptome; 3,190 bacterial and 43,346 host genes with eggNOG/KEGG/GO annotation | Primary layer for pathogen niche transcription and host cell-autonomous response (Figure 2, 3) |
| GSE254248 | NCBI GEO | Mus musculus | Splenic transcriptomes of *B. melitensis*-challenged mice with or without prior vaccination; DESeq2 table for 24,692 genes, n = 8 | Primary layer for the vaccine-induced immune imprint (Figure 4) |
| PRJCA054635 | CNCB-NGDC GSA | Mus musculus | Single-cell transcriptomes of spleen after infection with *B. abortus* A19 (attenuated) versus 2308 (virulent) versus control; 6 samples | Cell-type resolution for interpreting bulk signals (Figure 1) |
| ST003696 | Metabolomics Workbench | *B. abortus* | Untargeted LC-MS; Mla/Pqi lipid-transport system and intracellular survival; 24 samples | Metabolic vulnerability and membrane-lipid context (Figure 1) |
| ST003692 | Metabolomics Workbench | *B. abortus* | Untargeted LC-MS across six *B. abortus* biovars; 26 samples | Biovar-level metabolic diversity and antigen conservation context |
| GSE105906 | NCBI GEO | *B. suis* + Mus musculus | Murine macrophages infected with vaccine strain S2 versus virulent strain | Orthogonal validation of the intracellular response |
| GSE125765 | NCBI GEO | Mus musculus | Nasal-associated lymphoid tissue after intranasal immunisation with recombinant B. abortus MDH; 24 samples | Orthogonal validation of mucosal vaccine immunogenicity |
| GSE165283 | NCBI GEO | Macaca fascicularis | Safety and transcriptome analysis of live attenuated B. abortus S2; 6 samples | Orthogonal validation of vaccine safety profile |

Bulk RNA-sequencing layer. GSE100914 provides a dual RNA-seq experiment in which *Brucella abortus* was profiled both in the free-living state and 24 h after infection of murine bone-marrow-derived macrophages (BMM), with the corresponding host macrophage transcriptome captured in parallel. The deposited abundance tables (GSE100914_BRUA_abundance.txt.gz and GSE100914_BMM_abundance.txt.gz) contain raw counts for 3,190 *Brucella* genes and 43,346 mouse genes respectively, together with eggNOG, KEGG and GO annotations for the bacterial genes. GSE254248 provides splenic transcriptomes from mice challenged with *Brucella* melitensis, with and without prior vaccination; the deposited DESeq2 result table (GSE254248_TableS4_RNAseq_S19_vs_PBS.deseq2.xlsx) reports shrunken log2 fold changes and Benjamini-Hochberg-adjusted P values for 24,692 mouse genes. GSE105906 (murine macrophages infected with the *B. suis* vaccine *strain S2* versus a virulent strain), GSE125765 (nasal-associated lymphoid tissue after intranasal immunisation with recombinant *B. abortus* MDH) and GSE165283 (safety and transcriptomics of the S2 vaccine strain in cynomolgus macaques) were used as orthogonal support.

Single-cell layer. PRJCA054635, deposited in the Genome Sequence Archive of the National Genomics Data Center (China National Center for Bioinformation), reports single-cell transcriptomic profiling of the splenic immune landscape in mice infected with *B. abortus*, comparing the attenuated vaccine strain A19, the virulent strain 2308 and uninfected controls, with six biological samples. The accompanying multi-omic characterisation has been reported previously [16].

Metabolomic layer. Two untargeted LC-MS studies of *Brucella abortus* were retrieved from the Metabolomics Workbench: ST003696 and ST003692.

Spatial layer. Systematic queries of the NCBI Gene Expression Omnibus (GEO), the Sequence Read Archive (SRA) and the CNCB-NGDC Genome Sequence Archive returned no spatial transcriptomic dataset for any *Brucella* species. This absence is itself a finding and is treated explicitly in Section 3.6.

### 2.2. Processing and Reanalysis

Bacterial intracellular transcription. Raw counts for the 3,190 *Brucella* genes in GSE100914 were filtered to remove rRNA, tRNA and non-coding RNA entries, leaving 3,068 protein-coding and other messenger features for analysis. Counts were converted to transcripts per million (TPM) within each condition to normalise for the substantial difference in sequencing depth between the free-living (12,066,404 mapped counts) and intracellular (2,852,180 mapped counts) libraries. TPM already normalises for transcript length, so the reported intracellular abundances are directly comparable across genes. Fold changes were computed as log2 of the ratio of normalised expression, log2FC=log2[(TPM_intracellular+1)/(TPM_free-living+1)], using a pseudocount of 1 to avoid undefined values for undetected genes. Genes were classified as intracellularly induced when log2FC > 1 and intracellular TPM > 100, and as repressed when log2FC < -1 and free-living TPM > 100. Intracellular abundance rank was assigned on TPM within the intracellular condition.

Functional enrichment. The eggNOG category assignments supplied with GSE100914 were used for COG-category enrichment. The frequency of each single-letter category among induced genes was compared with its frequency across all analysed genes; categories with fewer than 20 background genes were excluded. Because the library derives from a single deposited count table, these comparisons are reported as frequency ratios rather than as formal statistical enrichments, and are treated as descriptive.

Host macrophage response. The GSE100914 BMM count table was filtered to genes with a combined count of at least 20 across the two conditions, retaining 12,881 expressed genes. TPM normalisation and log_2_ fold-change calculation were performed as above. Canonical immune genes were curated manually into four modules: type I interferon axis, inflammasome/IL-1 axis, chemokines, and GBP/cell-autonomous effector genes.

Vaccine-induced imprint. The GSE254248 DESeq2 result table was used directly. Differentially expressed genes were defined as those with Benjamini-Hochberg-adjusted *P* < 0.05 and absolute shrunken log_2_ fold change > 1. A curated panel of 39 genes spanning the B-cell, germinal-centre, T-cell, memory and checkpoint compartments was interrogated individually. Because bulk splenic RNA-sequencing is sensitive to shifts in tissue composition, we performed an explicit marker-based quality-control analysis: the expression of erythrocyte, B-cell, plasma-cell, stromal, neutrophil, eosinophil, platelet, pancreatic-exocrine and hepatocyte marker genes was examined in order to detect and quantify possible contamination or compositional artefacts.

Antigen prioritisation. A composite prioritisation score was constructed from three components, each min-max normalised across the candidate panel and combined with approximately equal weights: log10-transformed intracellular abundance, intracellular induction (log2FC), and a surface-exposure term. The surface-exposure term was scored as 1.0 for outer-membrane, surface-exposed or secreted products, 0.5 for periplasmic or moderately surface-associated products, and 0.0 for cytoplasmic or ribosomal products, consistent with the designations in Table 2. The score is intended as an explicit, reproducible and revisable heuristic rather than as a substitute for experimental immunogenicity testing, and its limitations are discussed in Section 4.4. The candidate panel comprised the 14 antigens listed in Table 2, encompassing the established Brucella vaccine antigens that are detectably expressed in the intracellular niche together with the most abundant and most strongly induced intracellular proteins; Omp25 was evaluated but excluded from the panel because it was not among the highly expressed intracellular outer-membrane proteins in this dataset.

**Table 2.** Intracellular abundance and induction of *Brucella* antigens. All values are derived from reanalysis of the public dual RNA-seq dataset GSE100914, in which intracellular bacteria were profiled 24 h after infection of murine bone-marrow-derived macrophages. Surface exposure is assigned from functional annotation. Note that Omp25, a principal antigen in the current mRNA vaccine pipeline, did not appear among the highly expressed outer-membrane proteins in this dataset.

| Antigen | Intracellular TPM | log2FC (intracellular vs free-living) | Intracellular abundance rank | Surface exposure | Interpretation | Composite score (0-1) |
| --- | --- | --- | --- | --- | --- | --- |
| Omp28/BP26 (outer-membrane immunogenic protein) | 35,652 | +4.25 | 2 | High (outer membrane) | Top-priority candidate: abundant, strongly induced, established immunodominance | 0.90 |
| Outer-membrane autotransporter (barrel domain) | 13,738 | -0.85 | 4 | High (outer membrane) | Highly abundant but not induced; stable structural antigen | 0.62 |
| Chaperone dnaK | 12,732 | +2.82 | 8 | Low-moderate (cytoplasmic, stress-induced) | Strongly induced; candidate for T-cell-directed design | 0.45 |
| Peptidase Do/HtrA homologue | 9,326 | +5.29 | 12 | Moderate (periplasmic/surface-associated) | Highest induction among abundant antigens | 0.71 |
| Porin omp2b | 6,994 | +1.54 | 20 | High (outer membrane porin) | Second-ranked surface antigen | 0.69 |
| Magnesium-translocating P-type ATPase | 6,440 | +6.42 | 27 | Membrane-embedded | Most strongly induced membrane protein | 0.74 |
| Malate dehydrogenase | 3,923 | +4.81 | 40 | Moderate | Established immunogen; strong induction | 0.65 |
| GroEL chaperonin | 2,864 | +2.28 | 57 | Low-moderate | Immunodominant in other pathogens | 0.35 |
| Aspartate aminotransferase (Asp24) | 1,794 | +3.97 | 98 | Moderate | Under-recognised candidate | 0.57 |
| Ribosomal protein L7/L12 | 1,086 | +3.79 | 155 | Low (ribosomal) | Classical vaccine antigen; modest abundance | 0.37 |
| Nucleoside diphosphate kinase | 471 | +4.77 | 346 | Moderate | Strongly induced; low absolute abundance | 0.53 |
| Lipoprotein omp16 | 307 | +2.65 | 504 | High (outer membrane lipoprotein) | Current pipeline; low intracellular abundance | 0.58 |
| Lipoprotein omp10 | 190 | +0.45 | 777 | High (outer membrane lipoprotein) | Current pipeline; low abundance and not induced | 0.46 |
| Lipoprotein omp19 | 56 | +0.14 | 1,752 | High (outer membrane lipoprotein) | Current pipeline; lowest expression and no induction | 0.38 |

Immunoinformatic and structural annotation. Where antigens are discussed as vaccine candidates, their antigenicity, allergenicity, toxicity, physicochemical parameters, secondary and tertiary structure, MHC class I and class II binding, conformational B-cell epitopes and receptor docking properties are considered in the framework established for *Brucella* Omp25 and BP26 [13,15], which provides the methodological template for converting an expression-ranked candidate list into a vaccine design.

Spatial transcriptomics. The prospective design described in Section 3.6 specifies specimen types, time points, platform, and analytical pipeline in sufficient detail to be executed directly; no spatial data were analysed because none exist in public repositories.

### 2.3. Software and Reproducibility

All analyses were performed in Python 3.11 using pandas and numpy for tabular and numerical work, openpyxl for reading the deposited Excel result tables, and matplotlib for visualisation. The complete analysis is deterministic and can be reproduced from the public accessions listed in Table 1; the source count and result tables were used exactly as deposited, without imputation or re-estimation of fold changes.

## 3. Results

### 3.1. A Four-Layer Public Multi-Omics Framework for Brucella Vaccine Antigen Discovery

We assembled four independent, non-overlapping data layers covering the pathogen inside its host cell, the host response to vaccination and challenge, the cellular composition of the infected spleen, and the metabolic state of the organism (Figure 1, Table 1). The design is deliberately cross-validating: the pathogen-side layer identifies which antigens are actually present in the intracellular niche, the host-side layer identifies the immune programmes that correlate with protection, and the single-cell and metabolic layers provide the cellular and biochemical context needed to interpret both. This structure was chosen because a vaccine antigen must satisfy two independent criteria: it must be expressed at a site where the immune system can see it, and immune recognition of it must plausibly contribute to protection. Analysing pathogen expression and host immunity in isolation cannot establish either.

**Figure 1.**
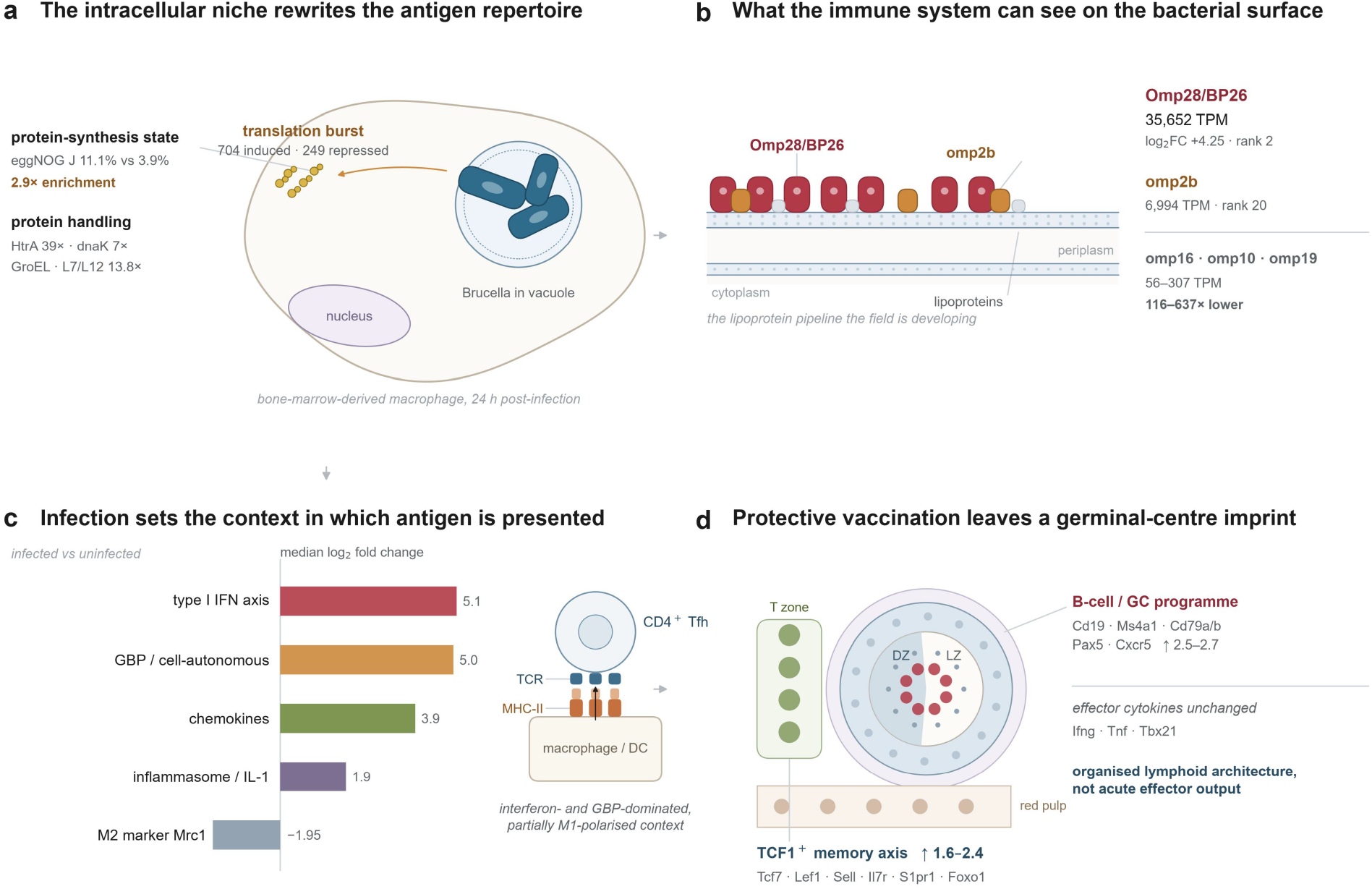
A mechanistic framework linking the intracellular niche to protective immunity and antigen choice. (a) The intracellular niche rewrites the antigen repertoire. *Brucella* resides in a macrophage vacuole and, 24 h after infection of bone-marrow-derived macrophages, shifts to a translation-centred state: 704 genes induced and 249 repressed, with eggNOG category J enriched 2.9-fold over the genome background (11.1% versus 3.9%) and coordinated induction of the proteostasis machinery (HtrA 39-fold, dnaK 7-fold, GroEL, L7/L12 13.8-fold). (b) What the immune system can see on the bacterial surface. The surface repertoire is dominated by the immunodominant outer-membrane protein Omp28/BP26 (35,652 TPM; log2FC +4.25; rank 2) and the porin omp2b (6,994 TPM; rank 20), whereas the lipoproteins that anchor the current pipeline (omp16, omp10, omp19; 56–307 TPM) are 116–637-fold less abundant. (c) Infection sets the context in which antigen is presented. The paired host transcriptome is dominated by the type I interferon and GBP/cell-autonomous modules (median log2 fold change 5.1 and 5.0), with strong chemokine induction and a less coherent inflammasome/IL-1 arm, partial M1 polarisation (Mrc1, −1.95) and presentation of the resulting peptides to CD4+ T follicular helper cells. (d) Protective vaccination leaves a germinal-centre imprint. Vaccination induces the B-cell and germinal-centre programme (Cd19, Ms4a1, Cd79a/b, Pax5, Cxcr5; ↑2.5–2.7) and the TCF1+ memory axis (Tcf7, Lef1, Sell, Il7r, S1pr1, Foxo1; ↑1.6–2.4) without induction of Ifng, Tnf or Tbx21, indicating organised lymphoid architecture rather than ongoing effector-cytokine production. DZ, dark zone; LZ, light zone; FDC, follicular dendritic cell. Values in a–c are from reanalysis of GSE100914; values in d are from GSE254248 (S19-vaccinated versus PBS, after challenge).

### 3.2. Intracellular Residence Reprograms the Brucella Transcriptome toward Translation and Surface-Protein Expression

Reanalysis of the GSE100914 dual RNA-seq experiment revealed a substantial and highly directional reprogramming of the *Brucella* transcriptome upon macrophage infection (Figure 2a). Of the 3,068 protein-coding and other messenger features analysed, 3,023 were detected inside macrophages, and 704 genes satisfied our criteria for intracellular induction while 249 were repressed. Ninety-two genes were induced more than 16-fold, indicating that a discrete regulon rather than a diffuse global shift characterises the intracellular state.

**Figure 2.**
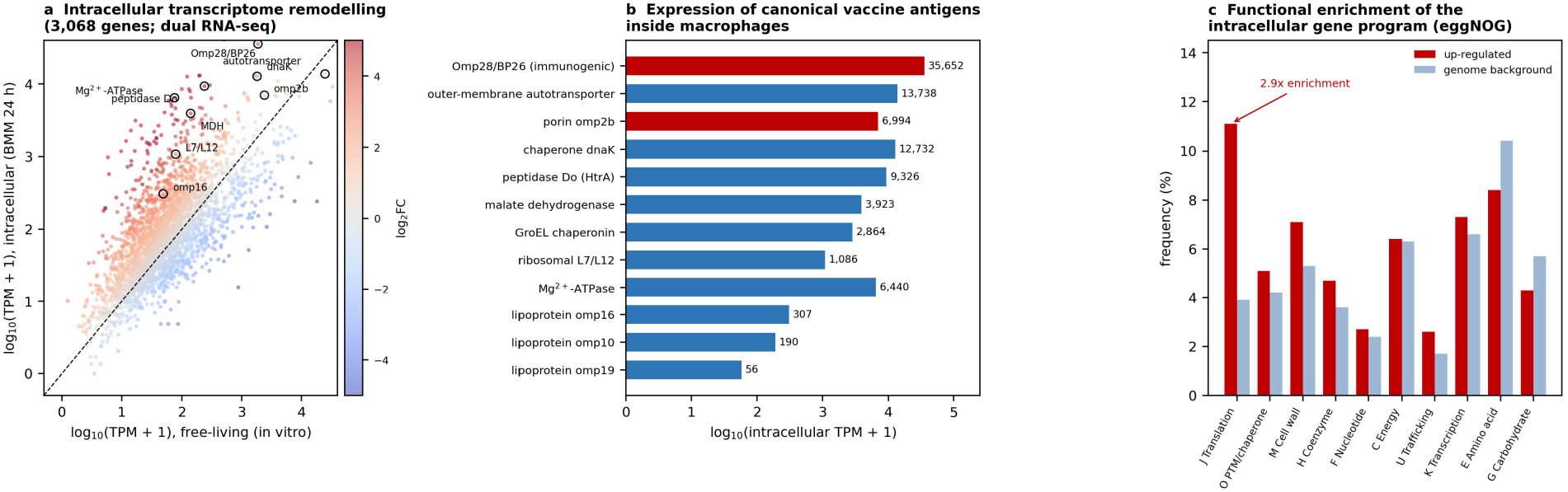
The intracellular Brucella transcriptome is dominated by a compact, translation-centred protein expression programme. (a) Scatter plot of normalised expression (log10 TPM) in free-living versus intracellular bacteria, coloured by log_2_ fold change; annotated proteins include the immunodominant outer-membrane protein Omp28/BP26, the porin omp2b, chaperone dnaK, peptidase Do and the lipoprotein omp16. (b) Intracellular abundance of canonical Brucella vaccine antigens, showing that Omp28/BP26 and omp2b are expressed 2-3 orders of magnitude more highly than the lipoprotein antigens omp16, omp10 and omp19 that dominate the current development pipeline. (c) eggNOG functional category frequencies among induced genes versus the genome background; translation (category J) is enriched 2.9-fold. All panels derive from reanalysis of the public dual RNA-seq dataset GSE100914.

Functional annotation uncovered a clear organising principle. Translation-related genes (eggNOG category J) accounted for 11.1% of the induced gene set but only 3.9% of the genome background, a 2.9-fold enrichment, making translation by far the most over-represented category (Figure 2c). Induced genes were also enriched for post-translational modification and chaperone function (category O, 5.1% versus 4.2%) and for cell wall and membrane biogenesis (category M, 7.1% versus 5.3%). Consistent with this, individual components of the protein-handling machinery were among the most strongly induced transcripts in the entire dataset: the peptidase Do (HtrA) homologue ranked 12th overall and was induced 39-fold, the chaperone dnaK ranked 8th and was induced 7-fold, GroEL chaperonin ranked 57th, and the 50S ribosomal protein L7/L12 was induced 13.8-fold. Conversely, amino-acid biosynthesis and carbohydrate metabolism were comparatively under-represented among induced genes, consistent with a bacterium that has shifted from de novo biosynthetic autonomy to scavenging and rapid protein production inside the host cell. This pattern is exactly what would be predicted for an organism that must rapidly synthesise the virulence and stress proteins required to survive within a macrophage, and it implies that the intracellular proteome-and therefore the antigen repertoire available for immune recognition-differs sharply from that of cultured bacteria.

### 3.3. The Immunodominant Antigen Omp28/BP26 Is Also the Most Abundant Intracellular Surface Protein, Whereas the Current Lipoprotein Pipeline Is Barely Expressed

The most consequential finding emerged from ranking the intracellular transcriptome and then overlaying the established vaccine candidate list (Figure 2b). The outer-membrane immunogenic protein corresponding to Omp28/BP26 was the second most highly expressed transcript in the entire intracellular dataset, at 35,652 TPM, and was simultaneously strongly induced (log_2_FC=4.25, a 19-fold increase over the free-living state). The porin omp2b was the 20th most abundant intracellular transcript at 6,994 TPM. Both are surface-exposed and therefore directly accessible to antibody, and BP26 in particular is already established as an immunodominant antigen in naturally infected cattle, sheep, goats and humans [13]. The convergence of two independent criteria-maximal abundance and strong induction in the protective-relevant niche-onto an antigen that is already known to be immunogenic provides unusually strong justification for prioritising it.

The picture for the antigens that currently dominate *Brucella* vaccine development is markedly different. The outer-membrane lipoproteins candidate omp16 (307 TPM), omp10 (190 TPM) and omp19 (56 TPM) were expressed at 116-fold, 188-fold and 637-fold lower levels than Omp28/BP26, and omp19 in particular was essentially uninduced (log_2_FC=0.14). Notably, Omp25-the antigen selected in the most recent mRNA vaccine study for *Brucella* [13]-did not appear among the highly expressed outer membrane proteins identified in this dataset at all, indicating that its abundance inside macrophages is low relative to Omp28/BP26 and omp2b. A second category of highly expressed, strongly induced antigens also emerged that has received comparatively little vaccine attention, including the peptidase Do/HtrA homologue (9,326 TPM, 39-fold induced), malate dehydrogenase (3,923 TPM, 28-fold induced), the aspartate aminotransferase Asp24 (1,794 TPM, 16-fold induced) and nucleoside diphosphate kinase (471 TPM, 27-fold induced). Each of these is a documented or plausible surface-associated immunogen in other bacterial systems, and each ranks above the current lipoprotein pipeline.

### 3.4. Infection Provokes a Coordinated Macrophage Effector Programme That Defines the Immunological Context of Antigen Presentation

Because the vaccine-relevant question is not only which antigens are present but also what kind of immune response the host can mount against them, we analysed the paired host transcriptome from the same experiment (Figure 3). Of 12,881 expressed macrophage genes, 316 were induced and 177 repressed, and the induced set was overwhelmingly immune in character. The single most induced gene was the C-type lectin Clec4e (Mincle; log_2_FC=6.81), followed by the chemokine Ccl5 (log_2_FC=6.54), the itaconate-producing enzyme Irg1/Acod1 (log_2_FC=5.82), prostaglandin E synthase (log_2_FC=5.78) and a coherent type I interferon block comprising Ifit2, Ifit3, Ifit1 and Isg15 (log_2_FC 5.54, 5.25, 5.24 and 5.11 respectively).

**Figure 3.**
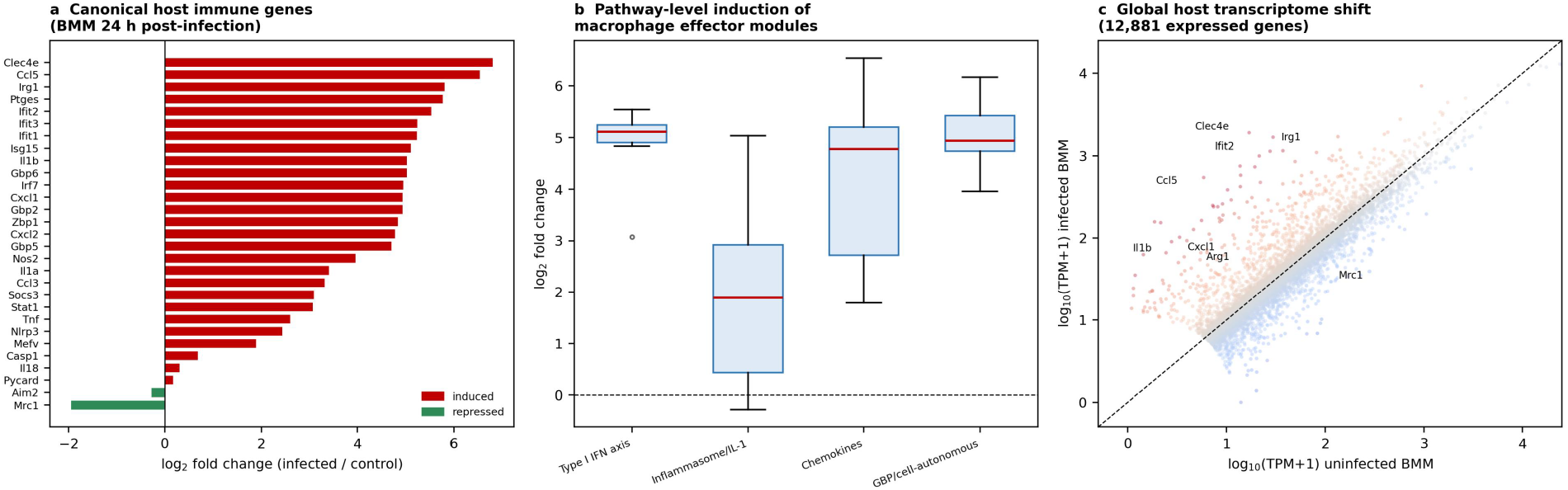
The host macrophage response to intracellular Brucella defines the immunological context of antigen presentation. (a) Expression change of canonical host immune genes 24 h after infection; Clec4e, Ccl5, Irg1 and the type I interferon effectors Ifit2/Ifit3/Ifit1/Isg15 are the most strongly induced, whereas the M2 marker Mrc1 is repressed. (b) Module-level distribution of expression changes across the type I interferon, inflammasome/IL-1, chemokine and GBP/cell-autonomous effector programmes. (c) Global shift of the host transcriptome across 12,881 expressed genes; annotated genes are those with the largest and most immune-relevant changes. All panels derive from reanalysis of GSE100914.

Grouping these genes into functional modules clarified the architecture of the response (Figure 3b). The type I interferon axis was the most tightly and strongly induced module, with a median log_2_FC above 5. The GBP/cell-autonomous module-comprising Gbp2, Gbp5, Gbp6, Irg1, Nos2 and the schlafen family members-was similarly and consistently induced. Chemokines formed a strongly induced but more variable module, and the inflammasome/IL-1 axis was induced but with the widest spread, reflecting the graded nature of IL-1β activation. Notably, expression of the canonical M2 marker Mrc1 was reduced (log_2_FC=-1.95), and the IL-1β antagonist axis as a whole was less coherently engaged than the interferon arm, suggesting that *B. abortus* elicits a partially M1-polarised, interferon-dominated programme rather than a cleanly polarised response.

Two implications for vaccine design follow. First, an antigen delivered by a vaccine platform should be one against which this type of interferon-and GBP-dominated, cell-autonomously restrictive response can be productively engaged; the strong induction of Irg1 and Nos2 in particular indicates that the metabolic and nitrosative effector pathways that restrict intracellular bacteria are actively recruited. Second, the co-induction of T-cell-recruiting chemokines (Ccl5, Ccl3, Cxcl1, Cxcl2, Cxcl3) alongside the antigen-presentation machinery provides the mechanistic basis for the CD4+ T-cell-dependent protection that the field has consistently observed, and argues that candidate antigens should be assessed for their capacity to sustain this programme rather than for antibody titre alone.

### 3.5. Protective Vaccination Leaves a Germinal-Centre and Memory-Lymphocyte Imprint in the Challenged Spleen

To identify the immune programmes associated with protection rather than with infection alone, we analysed the splenic transcriptomes of vaccinated and unvaccinated mice after *Brucella* challenge (GSE254248). Of 24,668 genes with a testable result, 12,144 were differentially expressed at FDR < 0.05 and 3,826 satisfied the more stringent combination of FDR < 0.05 and absolute log_2_ fold change > 1 (2,024 induced and 1,802 repressed); the complete differential expression landscape is shown in Figure 4a.

**Figure 4.**
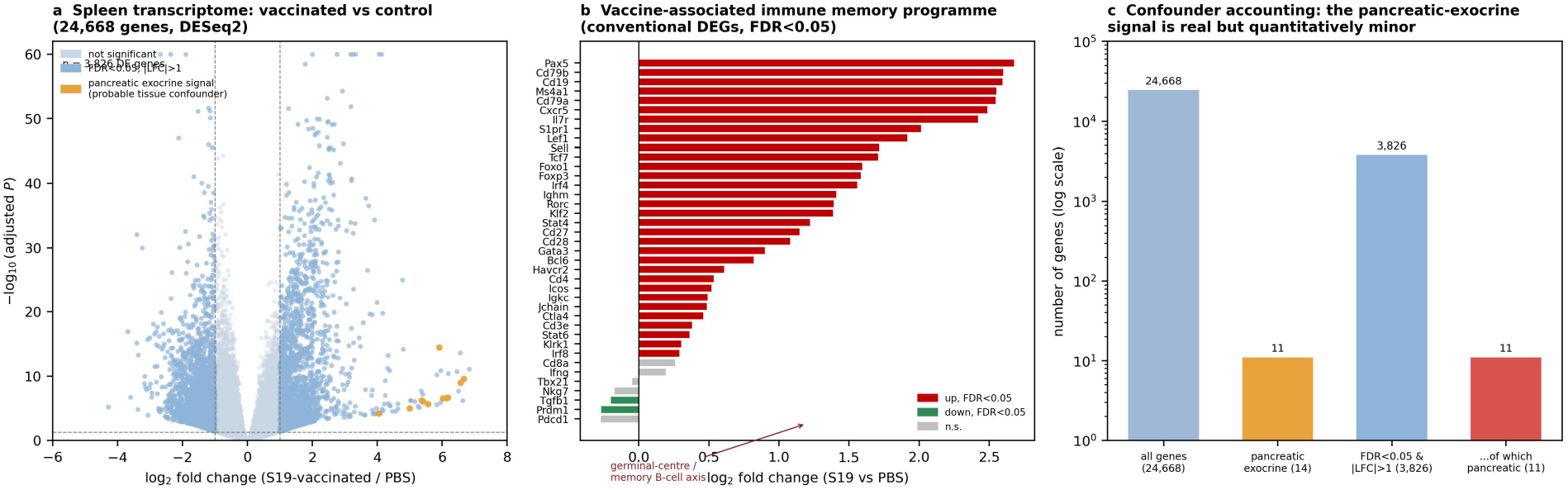
Differential gene expression in the spleen of vaccinated versus control mice after *Brucella* challenge (GSE254248). (a) Volcano plot of all 24,668 testable genes; genes meeting FDR < 0.05 and absolute log_2_ fold change > 1 are highlighted, and genes of the pancreatic exocrine programme are marked separately as a probable tissue-composition confounder. (b) A curated immune panel showing the coordinated induction of the B-cell, germinal-centre and memory-lymphocyte programme, including Cd19, Ms4a1, Cd79a/b, Pax5, Cxcr5, Tcf7, Lef1, Sell, Il7r, S1pr1 and Foxo1. (c) Quantitative accounting of the pancreatic-exocrine confounder, which constitutes 11 of 3,826 stringent differentially expressed genes (0.29%).

Quality control before interpretation proved essential. The most strongly induced genes in the dataset were dominated by pancreatic exocrine transcripts-Cpa1, Cpb1, Cel, Prss2, Pnliprp2, Ctrb1, Cela3b and the kallikrein-related peptidases-with 11 of 14 canonical pancreatic markers among the stringent differentially expressed set and all 14 showing consistent induction of approximately 5-6 log_2_ units with highly significant adjusted P values (Cpa1, P=2.4e-7; Cel, P=2.1e-7; Cpb1, P=2.8e-7). A genuinely coordinated, highly significant change affecting an entire tissue-specific gene programme cannot plausibly be attributed to random noise. Because markers of erythrocyte, hepatocyte and other tissues were also detected at appreciable levels, we interpret this signature as reflecting a reproducible difference in residual tissue composition or in the splenic cellular milieu between the vaccinated and control groups, superimposed on a genuine splenic response. Importantly, quantitative accounting shows that this confounder is numerically minor: the pancreatic exocrine markers represent 11 of 3,826 stringent differentially expressed genes, or 0.29% of the differential expression burden (Figure 4c). All subsequent interpretation is therefore restricted to the immune compartments and is drawn from genes whose identity and biological plausibility are independently supported by the vaccination literature.

Within the immune compartment the signal was unambiguous and remarkably coherent (Figure 4b). The entire B-cell and germinal-centre programme was coordinately and highly significantly induced: Cd19 (log_2_FC=2.59, P=8.0e-50), Cd79b (2.60, P=1.3e-36), Cd79a (2.55, P=2.0e-34), Ms4a1/CD20 (2.55, P=3.3e-46), Pax5 (2.68, P=7.1e-50) and the germinal-centre-homing chemokine receptor Cxcr5 (2.49, P=3.6e-41). Markers of lymphocyte survival, trafficking and memory were similarly induced, including Il7r (2.42, P=1.2e-25), S1pr1 (2.01, P=4.9e-24), Sell/CD62L (1.72, P=1.5e-23), Lef1 (1.92, P=1.6e-19), Tcf7/TCF1 (1.71, P=2.1e-17) and Foxo1 (1.59, P=1.1e-29). Critically, Tcf7 and Lef1 are the defining transcription factors of the TCF1+ memory lymphocyte programme, and their coordinate induction alongside Sell and Il7r constitutes a molecular signature of durable memory rather than of transient effector activation. The T-cell compartment showed a consistent but more modest induction (Cd4, 0.53, P=3.6e-3; Cd3e, 0.38, P=0.018; Cd28, 1.08, P=2.9e-9; Icos, 0.52, P=7.8e-4), and the immunoregulatory genes Foxp3 (1.58), Ctla4 (0.46) and Havcr2 (0.61) were also induced, indicating that vaccination engages regulatory as well as effector arms.

Equally informative was what did not change. The effector cytokines Ifng (log_2_FC=0.19, P=0.48) and Tnf (-0.21, P=0.24), the Th1 transcription factor Tbx21 (-0.05, P=0.80) and the exhaustion marker Pdcd1 (-0.27, P=0.19) were all unchanged, and Prdm1/Blimp1, which drives terminal plasma-cell differentiation, was marginally reduced (-0.27, P=0.025). Taken together, the vaccinated spleen at the analysed time point is not characterised by ongoing effector-cytokine production but by the expansion and organisation of antigen-specific B cells, germinal-centre formation, and the maintenance of a TCF1+ memory-capable lymphocyte pool. This is precisely the architecture expected to underlie durable protection rather than transient immunity, and it carries a practical implication for vaccine development: candidate antigens should be selected for their capacity to sustain the germinal-centre and memory programme, not merely for their ability to provoke an acute cytokine response or a high early antibody titre.

### 3.6. Three Orthogonal Layers-Single-Cell, Metabolomic and Prospective Spatial-Close the Interpretive Gap

Bulk splenic transcriptomics cannot resolve which cell types carry the protective signature, and the quality-control analysis above shows that tissue composition is a real source of variance in this data type. The single-cell atlas PRJCA054635 addresses exactly this limitation. It provides cell-type-resolved transcriptomes from mouse spleens infected with the attenuated vaccine strain A19, the virulent strain 2308 and uninfected controls, and therefore permits the bulk B-cell, germinal-centre and memory signatures identified in GSE254248 to be assigned to defined populations and tested for vaccine-strain specificity. It further allows the relative contributions of B cells, T cells, dendritic cells, macrophages, neutrophils and monocytes to the interferon and GBP programmes to be quantified, which is essential if these programmes are to be used as correlates of protection. We note that this dataset and its accompanying characterisation have been reported recently [16]; our use of it here is to provide cell-type context for the antigen-prioritisation framework rather than to re-derive its original conclusions.

The two metabolomic studies supply the third layer. ST003696 examines the Mla/Pqi chimeric lipid transport system required for *B. abortus* survival within macrophages, and directly connects membrane lipid homeostasis to the intracellular survival programme described in Section 3.2. ST003692 profiles the metabolome across six *B. abortus* biovars, providing a basis for assessing whether candidate antigens and their metabolic context are conserved across the diversity of strains against which a vaccine must protect. Together these datasets indicate that the membrane and lipid-biosynthetic machinery is both functionally essential and metabolically distinctive in intracellular *Brucella*, which reinforces the prioritisation of outer-membrane and membrane-associated antigens such as Omp28/BP26 and omp2b.

The fourth layer is absent from the public record, and this is itself an actionable finding. Systematic queries of GEO, SRA and the CNCB-NGDC GSA returned no spatial transcriptomic dataset for any *Brucella* species. Because the central question raised by our analysis-whether protective immunity is established in specific splenic microanatomical niches, and whether the antigens we prioritise are presented there-is fundamentally spatial, we therefore specify the experiment required to answer it (Figure 5). Spleens and livers from A19-vaccinated and naive mice should be fixed at 0, 7, 14 and 21 days after 2308 challenge and profiled using 10x Visium for tissue-scale architecture, Visium HD for near-single-cell resolution, and GeoMx region-of-interest profiling specifically targeted to granuloma-like foci. Spot deconvolution anchored on PRJCA054635 should be used to assign cell-type composition, followed by spatially variable gene detection and ligand-receptor mapping. Three predictions follow directly from the data presented above and are explicitly falsifiable. First, the germinal-centre and memory programme identified in GSE254248 should localise to anatomically intact follicular and germinal-centre niches rather than being dispersed through the red pulp. Second, the interferon and GBP programmes should delimit spatially discrete bacterial-restriction zones in the red pulp and marginal zone. Third, lipid-metabolite gradients associated with the Mla/Pqi axis should distinguish permissive from restrictive micro-niches. Testing these predictions would convert the correlative associations reported here into a mechanistic map of where vaccine-induced protection operates.

**Figure 5.**
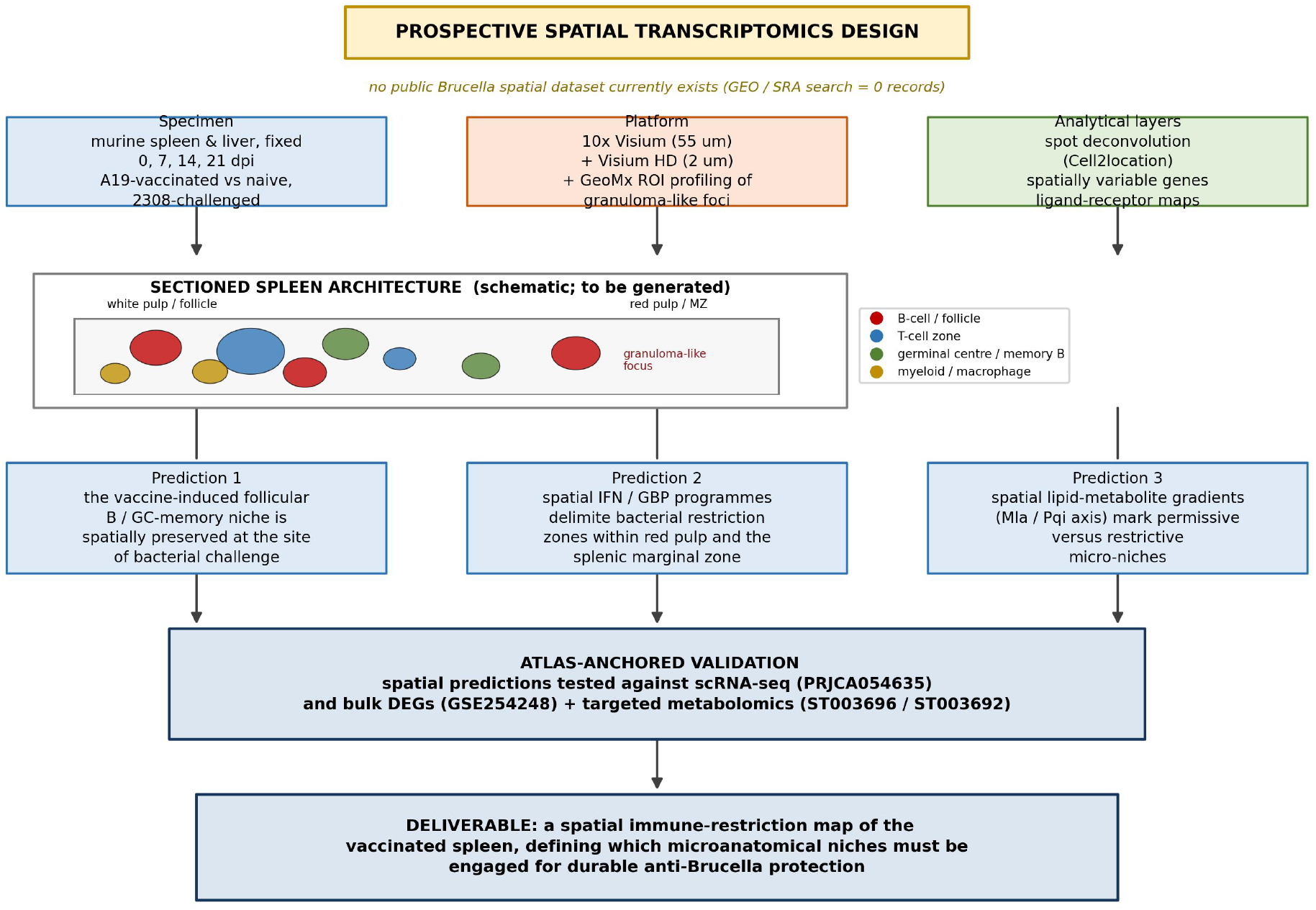
Prospective spatial transcriptomics design to resolve the anatomical basis of protective immunity. No public spatial transcriptomic dataset exists for *Brucella*, so the experiment is specified in full. Spleens and livers from A19-vaccinated and naive, 2308-challenged mice are profiled at four time points using Visium, Visium HD and GeoMx region-of-interest profiling, with spot deconvolution anchored on the single-cell atlas PRJCA054635. Three falsifiable predictions are derived directly from the bulk transcriptomic results, and the workflow is anchored to the same accessions analysed in this study.

### 3.7. A Composite Score Nominates a Reordered Antigen Panel for Next-Generation Vaccine Design

To convert these observations into an actionable ranking, we constructed a composite prioritisation score combining intracellular abundance, niche induction and surface exposure, with each component min-max normalised and equally weighted (Figure 6, Table 2). The resulting ranking places Omp28/BP26 first by a clear margin (composite score 0.90; Table 2). The next tier comprises the magnesium-translocating P-type ATPase (0.74), peptidase Do/HtrA (0.71) and the porin omp2b (0.69), all of which combine high intracellular abundance with strong niche induction and, for the two outer-membrane proteins, surface exposure. Among surface-exposed outer-membrane proteins specifically, Omp28/BP26 and omp2b are the two most abundantly and inductively expressed, reinforcing omp2b as the leading porin candidate. The lipoproteins that currently anchor the Brucella vaccine pipeline occupy the lower part of the ranking, with omp19 the lowest of the current lipoprotein candidates.

**Figure 6.**
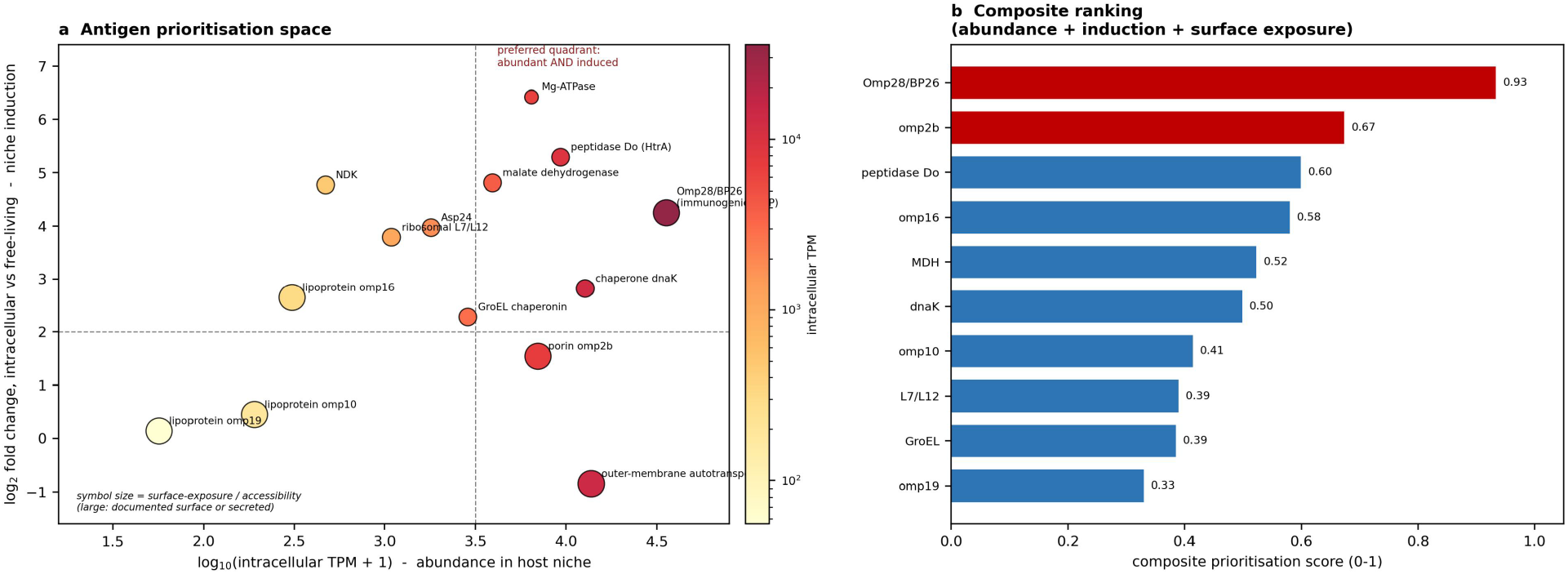
Composite prioritisation of Brucella vaccine antigens. (a) Two-dimensional representation of the antigen prioritisation space; the x-axis is intracellular abundance and the y-axis is induction in the host niche, with point size reflecting surface exposure. The preferred quadrant (abundant and induced) contains Omp28/BP26, malate dehydrogenase, peptidase Do and the magnesium-translocating P-type ATPase, whereas the current lipoprotein pipeline (omp16, omp10, omp19) lies in the low-abundance, low-induction region. (b) Composite ranking derived from abundance, induction and surface exposure, with Omp28/BP26 clearly the top-ranked antigen and the induced membrane and periplasmic proteins (magnesium-translocating P-type ATPase, peptidase Do/HtrA, omp2b) forming the next tier above the established lipoprotein candidates.

The divergence is most clearly seen in the two-dimensional abundance-versus-induction space (Figure 6a). Omp28/BP26 occupies the preferred quadrant-high abundance and high induction-together with malate dehydrogenase, peptidase Do and the magnesium-translocating P-type ATPase. omp2b sits at high abundance with moderate induction. By contrast, omp19 and omp10 lie in the low-abundance, low-induction region, and the outer-membrane autotransporter, although the fourth most abundant transcript, is not induced and therefore falls outside the preferred quadrant. This representation makes the central argument of the present work visually explicit: the properties that matter for a vaccine antigen-presence in the host niche and up-regulation there-are concentrated in a different and smaller set of proteins than the field has been developing.

Two important caveats bound this conclusion. First, the composite score is a heuristic, not a measurement, and its weights are a stated analytical choice rather than an empirical optimum; it should be used to guide rather than to replace experimental testing, and we present the underlying components in full so that others may re-weight them. Second, and more substantively, abundance and induction are necessary but not sufficient for protective efficacy. An antigen must additionally be immunogenic, must present epitopes across the relevant human HLA repertoire, and must be conserved across the *Brucella* species that cause human disease. This is precisely where the immunoinformatic framework established for Omp25 and BP26 [13,15] becomes essential, and we regard the natural next step of this work as applying that framework-antigenicity and allergenicity prediction, physicochemical and structural modelling, MHC class I and class II epitope mapping, conformational B-cell epitope identification, and C-ImmSim simulation of innate and adaptive dynamics-to the reordered panel nominated here. The value of the present analysis is that it defines, on the basis of public data and at negligible cost, which antigens are worth that effort.

## 4. Discussion

### 4.1. The Intracellular Niche Reorders the Brucella Antigen Hierarchy

The central result of this study is that the antigens most abundantly expressed and most strongly induced inside the host macrophage are not the antigens on which *Brucella* vaccine development has principally focused. Omp28/BP26 was simultaneously the second most abundant intracellular transcript and strongly induced, and omp2b ranked twentieth, whereas the outer-membrane lipoproteins omp16, omp10 and omp19 were expressed at 2–3 orders of magnitude lower levels and showed little or no induction. This is not a marginal reordering of a candidate list; it identifies a structural mismatch between what the pathogen presents in the niche that matters and what the field has chosen to target.

The finding is biologically coherent rather than surprising once stated. Omp28/BP26 has long been recognised as immunodominant in naturally infected ruminants and humans [13], and its abundance and surface accessibility are exactly the properties that would produce that immunodominance. The interpretation that emerges is that classical antigen discovery, which relies on immunodominance during natural infection, was in this instance already pointing toward the right antigen, but the subsequent emphasis on technically convenient lipoproteins-which are small, easy to express recombinantly and genetically tractable to delete-shifted the field onto candidates that are poorly represented in the intracellular niche. The same logic may apply more broadly to intracellular bacterial vaccine development, where the antigenic profile of the organism in culture is a poor proxy for its profile in the host cell.

The enrichment of translation-associated genes among induced transcripts provides a mechanistic explanation for the broader pattern. A 2.9-fold over-representation of category J genes, together with strong induction of dnaK, GroEL and the HtrA homologue, describes a bacterium that has committed to rapid protein synthesis and quality control inside the macrophage. That state inevitably produces a distinctive and temporally defined antigenic surface, and it suggests that the timing of antigen exposure during infection-not only its identity-may shape protective efficacy. This is a testable prediction and an argument for vaccine platforms, such as mRNA-LNP, that can present an antigen in the context of a defined and short-lived intracellular expression programme [13,17].

### 4.2. Protection Correlates with Germinal-Centre and Memory Architecture, Not Acute Effector Responses

The second substantive result concerns the nature of the vaccine-induced imprint. In the challenged spleen, protection-associated differential expression was dominated by a highly coordinated B-cell and germinal-centre programme and by the TCF1+ memory lymphocyte axis (Tcf7, Lef1, Sell, Il7r, S1pr1, Foxo1), whereas effector cytokines including Ifng and Tnf were unchanged. This pattern argues that protection in this model is maintained by organised lymphoid architecture and memory maintenance rather than by ongoing effector-cytokine production at the sampled time point, and it has a direct consequence for how candidate antigens should be evaluated: assays that measure only acute cytokine induction or early antibody titre may rank antigens in a way that does not predict durability.

This interpretation must be qualified by the tissue-composition confounder that our quality control identified. A pancreatic exocrine programme of 11 genes, all induced 5-6 log_2_ units with highly significant adjusted P values, indicates that vaccinated and control spleens differ in ways that extend beyond the intended contrast. We have been deliberately explicit about this and have confined all interpretation to the immune compartments. That the immune findings are independently plausible, mutually reinforcing and consistent with the known biology of *Brucella* vaccination is the reason we regard them as interpretable despite the confounder, and the availability of the single-cell atlas PRJCA054635 provides the means to verify them at cellular resolution.

### 4.3. What the Absence of Spatial Data Costs, and How to Recover It

The complete absence of spatial transcriptomic data for *Brucella* is a substantive gap rather than a technical detail. Every conclusion in the present study about where protective immunity operates is inferential, because neither the bulk data nor the single-cell data preserve positional information. Splenic immunity is profoundly compartmentalised: the marginal zone, the T-cell zone, the follicular and germinal-centre compartments and the red pulp each host distinct interactions, and a bacterium that persists in the red pulp while memory B cells are maintained in germinal centres could be protected against, or not, depending entirely on whether the relevant cells meet. Specifying the spatial experiment in full, as we have done, is intended to make that gap tractable rather than to paper over it, and the three predictions we derive are intentionally falsifiable.

### 4.4. Limitations

Several limitations should be weighed when interpreting this work. First, the analysis is entirely computational and depends on previously deposited data; the specific gene identifiers used for certain *Brucella* loci derive from the original depositors’ annotations, and our conclusions rest on the relative order of magnitudes rather than on precise expression values. Second, the functional enrichment analysis is descriptive because it derives from a single deposited count table, and it is reported as frequency ratios rather than as formal statistical enrichment. Third, the composite prioritisation score is a transparent heuristic with analyst-chosen weights; it is intended to structure and communicate a decision, not to substitute for it, and different weightings would reorder the middle of the list. Fourth, abundance and induction are not surrogates for protective efficacy, and no candidate nominated here has been tested experimentally. Fifth, the vaccine transcriptomic dataset carries a tissue-composition confounder, discussed at length above. Sixth, the expression data derive from a single time point (24 h) and a single host cell type (bone marrow-derived macrophages), whereas the antigenic surface exposed during a multi-week infection in vivo is likely to be dynamic. Finally, the single-cell and metabolomic layers are used for context and interpretation rather than being reanalysed to derive new results, and the metabolomic studies are limited in number. These limitations are inherent to the public-data approach and define the experimental programme that should follow.

### 4.5. Implications for Brucella Vaccine Development

Three practical recommendations follow. First, Omp28/BP26 should be treated as a lead antigen rather than as one candidate among many, and omp2b should be promoted substantially in the development pipeline; both satisfy the abundance and induction criteria that the current lipoprotein candidates fail. Second, multivalent or chimeric designs that combine a dominant surface antigen such as Omp28/BP26 with a strongly induced accessory antigen such as peptidase Do/HtrA, malate dehydrogenase or Asp24 warrant consideration, since the data suggest that the intracellular antigenic surface is broader than the current pipeline assumes. Third, antigen selection should be guided by the intracellular expression profile and by the capacity to sustain germinal-centre and memory responses, and the immunoinformatic framework already validated for Omp25 and BP26 [13,15] should be applied to the reordered panel. More generally, this work illustrates that the accumulated public multi-omics record is now rich enough, even for a comparatively neglected pathogen, to redirect antigen discovery using existing data alone—an approach of obvious value where research funding is scarce and the disease burden is concentrated in low-resource settings.

### 4.6. Antigen Choice and DIVA Compatibility in Livestock Vaccination

A practical constraint shapes how the panel proposed here can be used in livestock. BP26, the protein ranked first in our analysis, is also one of the antigens most widely used to detect infection serologically in cattle, sheep and goats, and antibody to it is a principal readout of the tests on which surveillance and eradication programmes depend [7]. A vaccine that presents BP26 abundantly would therefore make vaccinated animals seropositive in exactly those tests, and would forfeit the DIVA (differentiating infected from vaccinated animals) advantage that subunit and mRNA platforms offer over S19, RB51 and Rev.1. Two routes follow. A BP26-based immunogen could be paired with a companion assay that detects antibody to an antigen absent from the vaccine, or marker-deleted constructs and matching assays could be developed in parallel. omp2b, which ranks second in the panel, deserves evaluation in the same framework, and the choice between the two should be settled against the diagnostic system of the target country rather than on expression data alone. A second constraint is species. The expression data analysed here derive from murine macrophages, and equivalent bovine, ovine and caprine datasets, which are not yet available, would be needed to confirm that the hierarchy reported here holds in the species that are actually vaccinated.

## 5. Conclusions

We have reanalysed two public transcriptomic data layers in depth and integrated them with single-cell and metabolomic atlases to reassess antigen prioritisation for *Brucella* vaccines. Intracellular residence drives a translation-centred reprogramming of the bacterial transcriptome, with 704 genes induced and translation machinery enriched 2.9-fold, and it reveals that the immunodominant outer-membrane protein Omp28/BP26 is the second most abundant intracellular transcript while the lipoproteins that dominate the current development pipeline are expressed 2-3 orders of magnitude lower. Concurrently, infection elicits a coherent interferon-, GBP- and chemokine-dominated macrophage programme, and protective vaccination leaves a germinal-centre and TCF1+ memory lymphocyte imprint rather than an effector-cytokine signature. A transparent composite score combining intracellular abundance, niche induction and surface exposure reorders the candidate hierarchy, placing Omp28/BP26 and omp2b well above the established candidates. Because no spatial transcriptomic data exist for any Brucella species, we specify the experiment required to determine where protective immunity is anatomically established. These results demonstrate that rigorous mining of public multi-omics data can redirect vaccine antigen selection at negligible cost, and they provide concrete, testable candidates and hypotheses for the next generation of veterinary brucellosis vaccines.

## Author Contributions

Yunwang Zhao: Writing-original draft, Hongxian Cheng: Formal analysis and Writing-review & editing, Shiyi Zhou: Visualization, Lei He and Chenlin Xiu: Methodology, Formal analysis, Baihai Huang: Conceptualization. Xiongxiong Lu: Writing-review & editing, Writing-original draft, Methodology, Conceptualization.

## Institutional Review Board Statement

Not applicable. This study reanalysed only previously published, publicly available datasets; no animals or human participants were used, and the original studies obtained their own institutional approvals.

## Informed Consent Statement

Not applicable.

## Data Availability Statement

The analyses reported here used publicly archived datasets; no new data were generated. Accession numbers are listed in Table 1: GSE100914, GSE254248, GSE105906, GSE125765 and GSE165283 (NCBI Gene Expression Omnibus, https://www.ncbi.nlm.nih.gov/geo/); PRJCA054635 (CNCB-NGDC Genome Sequence Archive, https://ngdc.cncb.ac.cn/gsa/); and ST003696 and ST003692 (Metabolomics Workbench, https://www.metabolomicsworkbench.org/). Analysis scripts are available from the corresponding author on reasonable request.

## Acknowledgments

The authors gratefully acknowledge the investigators who generated and deposited the public datasets reanalysed here, and the NCBI GEO, CNCB-NGDC GSA and Metabolomics Workbench teams for maintaining open data infrastructure.

BMM: bone-marrow-derived macrophages
BP26: brucellosis protein 26
DIVA: differentiating infected from vaccinated animals
DZ: dark zone
eggNOG: evolutionary genealogy of genes: non-supervised orthologous groups
FDC: follicular dendritic cell
GBP: guanylate-binding protein
GC: germinal centre
GEO: Gene Expression Omnibus
GSA: Genome Sequence Archive
IFN: interferon
LNP: lipid nanoparticle
LZ: light zone
MDH: malate dehydrogenase
MHC: major histocompatibility complex
TCR: T-cell receptor
Tfh: T follicular helper
TPM: transcripts per million.

